# Robust brain inspired prediction of speech recognition accuracy across many words, voices, and natural background sounds

**DOI:** 10.64898/2026.04.27.721224

**Authors:** Alex C. Clonan, Ian H. Stevenson, Monty A. Escabí

**Affiliations:** Electrical and Computer Engineering, University of Connecticut, Storrs, CT 06269; Biomedical Engineering, University of Connecticut, Storrs, CT 06269; Psychological Sciences, University of Connecticut, Storrs, CT 06269; Institute of Brain and Cognitive Sciences, University of Connecticut, Storrs, CT 06269

## Abstract

Although humans generally excel at recognizing speech in noisy natural environments, performance varies widely due to differences in the background acoustics and individual talker attributes such as voice quality, intonation, and pitch. The underlying neural and acoustic factors of this perceptual variability remain elusive, making it an unresolved challenge to predict when speech recognition will succeed or fail. Here, we characterize human speech recognition across a wide range of natural conditions using speech spoken by 326 diverse male and female talkers of multiple ages embedded in 33 unique background noises. Across this vast soundscape, speech recognition is strongly driven by the spectrum and modulation statistics of the noise. Yet, articulatory speech features - including fundamental and formant frequencies - exert categorically distinct modulatory effects on recognition accuracy across the talker’s age and gender, and specific words. This perceptual variability is captured by a low-dimensional statistical representation of sounds based on computations that emulate intermediate stages of sound processing and feature extraction in auditory midbrain. A model utilizing these sound summary statistics accurately predicts individual listener’s single-trial recognition behavior, accounting for performance variance across individual voices, words, and backgrounds. Thus, speech-in-noise perception across diverse natural conditions relies largely on a simple set of spectrotemporal summary statistics known to be represented by central neural populations.

**Significance Statement:** Although humans can easily maintain conversations amidst environmental noise and voice variations, this capacity is severely degraded by hearing impairment and challenges modern speech recognition technologies. This study evaluates human speech recognition across a broad spectrum of environmental noises and diverse speaker demographics. We show that a brain-inspired auditory representation of sounds extracts critical statistical sound features driving human behavior, precisely predicting variations in perceptual accuracy across distinct voices, words, and environmental backgrounds.

## Introduction

Listening in the everyday world is seldom a simple task, since background sounds frequently distort or mask sounds of interest ^1^. Yet, the human auditory system is surprisingly robust to noise, allowing us to hear one another in a noisy restaurant filled with many voices or while walking down a city street with a cacophony of traffic and construction sounds. Recognizing speech in noise is a major challenge for individuals with hearing impairment, language processing differences, and even machine systems. Thus, understanding what speech and background factors impact the acuity, robustness, and efficiency of human hearing under natural distortions is of critical scientific and clinical interest. Recognition abilities vary based on the relative intensity of the foreground to the background, but even at a fixed signal-to-noise ratio (SNR), recognition accuracy varies widely across different backgrounds ^2,3^. The details of the speech – the specific talker, phonemes, their quality, intonation, and pitch – impact speech recognition as well ^4–7^, and these effects are likely background-dependent. Accurately accounting for the moment-by-moment contributions of the background, foreground, and interaction between them is an ongoing challenge for understanding the perception of speech in natural listening environments. Here, using a combination of human perceptual experiments and computational modeling, we aim to characterize foreground effects on speech recognition and foreground-background interactions on the timescale of single words.

Many previous studies have examined the specific sound features that impact speech perception, such as the fundamental frequency, consonants, phoneme-level features and other talker characteristics ^8–12^. *Energetic masking,* where the spectrum of the noise overlaps with or exceeds the energy of the competing speech, is a critical factor influencing speech perception ^13^. Since many environmental sounds are rich with transient acoustic events and are nonstationary, times of decreased power can allow speech to stand out, in a phenomenon known as glimpsing ^14,15^. *Informational* factors, such as spectrotemporal modulations that underlie word identity and contextual cues that influence word uncertainty ^13,16,17^, along with spatial cues ^18–21^ all also influence speech perception in noise. One limitation of many previous speech-in-noise studies is that they have not focused on the interactions between speech and natural background features and often have not focused on single-trial predictions. Here we aim to address these limitations by using highly diverse natural backgrounds and speech from multiple talkers to characterize the mechanisms underlying speech-in-noise perception.

Theories of auditory processing have proposed that humans rely on statistical sound information for perception and that statistics related to the spectral and modulation profiles of speech and stationary natural sounds can explain many aspects of human perception ^2,22,23^. Mid-level regions of the auditory pathway, such as the inferior colliculus, are specifically sensitive to both the spectral and modulation statistics of natural sounds ^24–26^ and neural activity in these subcortical regions can well describe both the identity of individual sound stimuli and categorical information about the sound type ^27–29^. Additionally, models based on spectrotemporal sound representations that account for modulation selectivity at this level can partly explain speech recognition abilities in natural noises ^2^.

Here we evaluate the hypothesis that spectrotemporal modulations in natural background sounds interfere with the representation of speech articulatory features in noise. Using both perceptual measurements and biologically plausible modelling techniques, we find that the ability to recognize speech varies widely depending on the foreground word, the speaker’s voice, and the background statistics. Some backgrounds give rise to consistent impairment in speech recognition, regardless of the foreground, while in some cases the background and foreground interact, and both determine listeners’ speech recognition accuracy. These interactions are linked to specific speech articulatory features, including fundamental and formant frequencies, which cluster across words as well as the talker’s age and gender. We find that the detailed behavioral effects can be directly explained on a sound-by-sound basis across words, talkers, and backgrounds by a model of central auditory representations of the foreground and background modulation statistics.

## Results

To determine how voice features and background statistics together impact speech recognition, we used a block randomized design and tested digit recognition abilities of n=10 participants using foreground English speech produced by many distinct speakers (n=326) mixed with many natural and perturbed background noises (9,900 mixtures; see Methods).

For a specific background, we mix foregrounds from different speakers producing the same word (Fig. 1A left) and the same speaker producing different words (Fig. 1A, right). By comparing across these trials, we can quantify how features of a speaker’s voice impact recognition in noise and how features of the word impact recognition in noise. By comparing recognition abilities in multiple backgrounds where the speaker (Fig. 1B left) or word (Fig 1B, right) is fixed, we can also determine to what extent foreground features interact with the spectral and temporal statistics of these different backgrounds. Participants were tasked with recognizing three digits (Fig. 1C) within natural background sounds (Fig. 1D, Original, OR; shown for the jackhammer background) and perturbed variants which isolate the sound spectrum (Fig. 1D, Spectrum Matched, SM) or the modulations (Fig. 1D, Modulation Matched, MM) and which allow determining how these statistical features influence speech perception ^2^. Sounds were block randomized across backgrounds and their perturbed variants and balanced across 4 different speaker populations (Adult Male, AM, n=111; Child Male, CM, n=50; Adult Female, AF, n=114; Child Female CF, n=51) (Fig. 1E). Altogether, participants’ recognition accuracy was measured across a wide range of natural conditions, allowing us to characterize and model the effects of different background statistics and foreground articulatory features.

**Figure 1:**
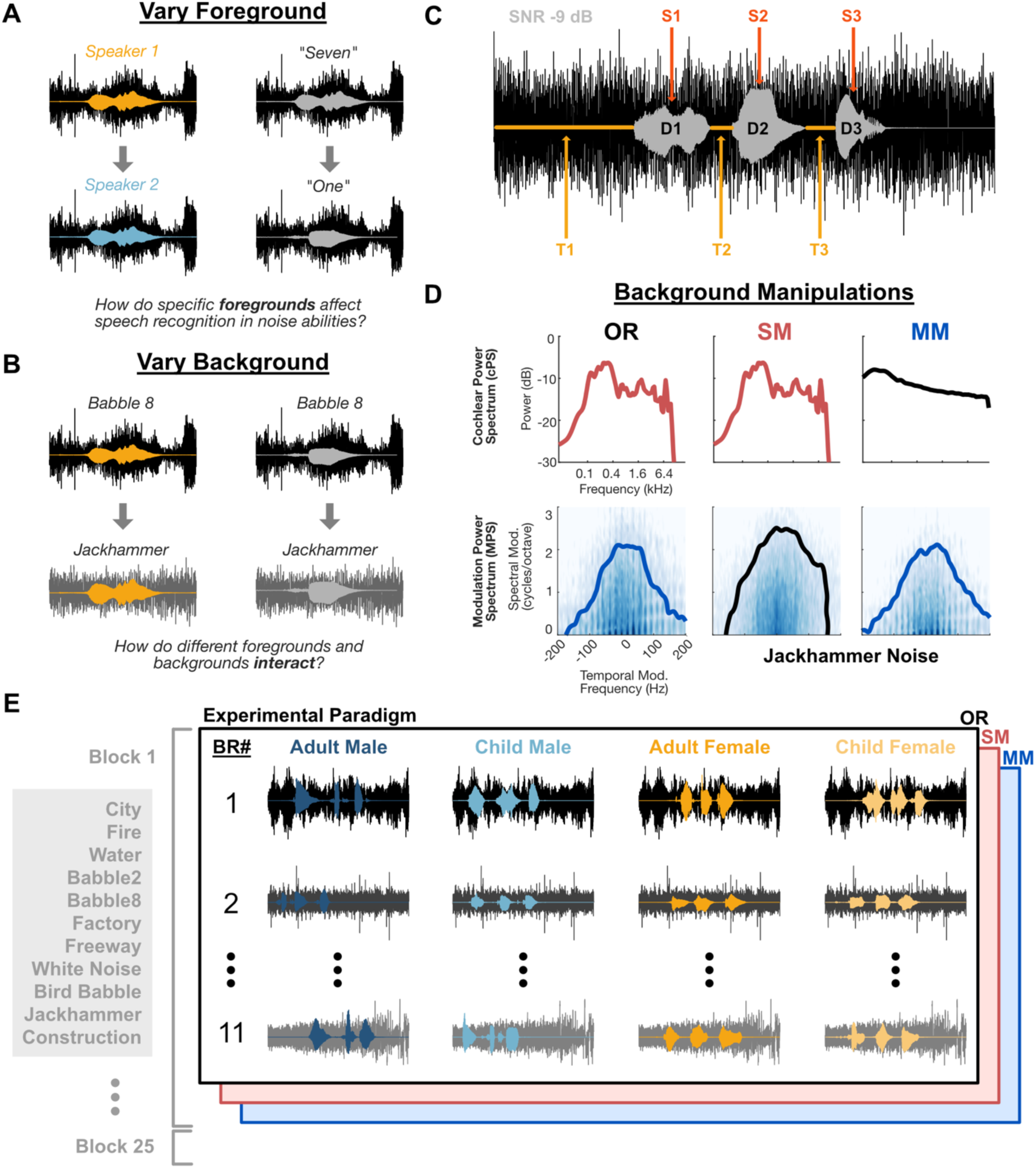
Measuring speech recognition sensitivity with variable foregrounds and backgrounds. To differentiate the effects of foreground voice and word-level acoustics we used a block randomized design consisting of trials that vary in foreground and background. A) For a specific background, the speaker (left, orange to blue) or the target word can vary (right, *seven* to *one*). B) To determine how specific speech features potentially interact with natural backgrounds statistics, we also compare speech recognition accuracy for the same voice (left) or target word (right), while varying the background. C) Speech in noise stimuli (SNR -9dB) were generated by mixing a background excerpt and three spoken digits (D1-D2-D3) from three variable speakers (S1-S2-S3) at random time intervals (T1-T2-T3). D) We tested original (OR) natural background sounds (e.g. fire, multi-speaker babble, water) along with white noise. Additionally, we generated perturbated backgrounds with matched spectral (SM) or modulation (MM) statistics. The cochlear power spectrum (cPS, top) and 90% power contour for the modulation power spectrum (MPS, bottom) are shown for an original (OR) excerpt of jackhammer (left), a spectrum matched (SM) perturbation (middle), and a modulation matched (MM) perturbation (right). E) In each experimental block, each of 11 backgrounds was presented for each background condition (OR, SM, and MM; totaling 33 backgrounds) and mixed with each of four voice groups: adult/child and male/female (AM, CM, AF, CF). Across different blocks (1-25), the specific exemplar of a given background changes.

### Accuracy variation is associated with foreground articulatory features

Participants listened to sounds over headphones (Fig. 2A) and were instructed to identify the digits in the sound mixture via keyboard entry (e.g., 7-9-2). In previous work, we found that digit recognition accuracy varied extensively across backgrounds depending on their spectrum and modulation statistics ^2^. Here we additionally find that accuracy varies based on the foreground speaker group (Fig. 2B1-3) and across digits (Fig. 2C1-3). Comparing the OR backgrounds, accuracy for babble-8 and factory backgrounds was the lowest, while accuracy for fire and white noise was the highest among the 11 backgrounds (see Methods). Additionally, by comparing accuracy, across OR, SM, and MM conditions, we can attribute performance to spectral or modulation statistics of the background. Across backgrounds, the voice group effects are influenced by the background spectrum and modulation cues, however they do not appear to be purely the result of one or the other. For babble-8 backgrounds, children’s speech is more recognizable than adult speech both for the OR background (15±2%, 36±3%, , AM, AF vs 59±2%, 63±3%, , CM, CF; mean±sem) and when the background modulations are whitened (SM, Fig. 2B2) (72±2%, 74±2%, , AM, AF vs 86±2%, , 85±1%, , CM, CF; mean±sem). This suggests that the background spectrum alone is sufficient to cause voice group effects. At the same time, even when the spectra are matched across backgrounds (MM, Fig. 2B3) the voice group effects still differ across backgrounds (15±2%, 34±3%, , AM, AF vs 52±2%, , 44±3%, , CM, CF; mean±sem), suggesting that the background modulations cues must also play a role in driving accuracy differences across voice groups.

**Figure 2:**
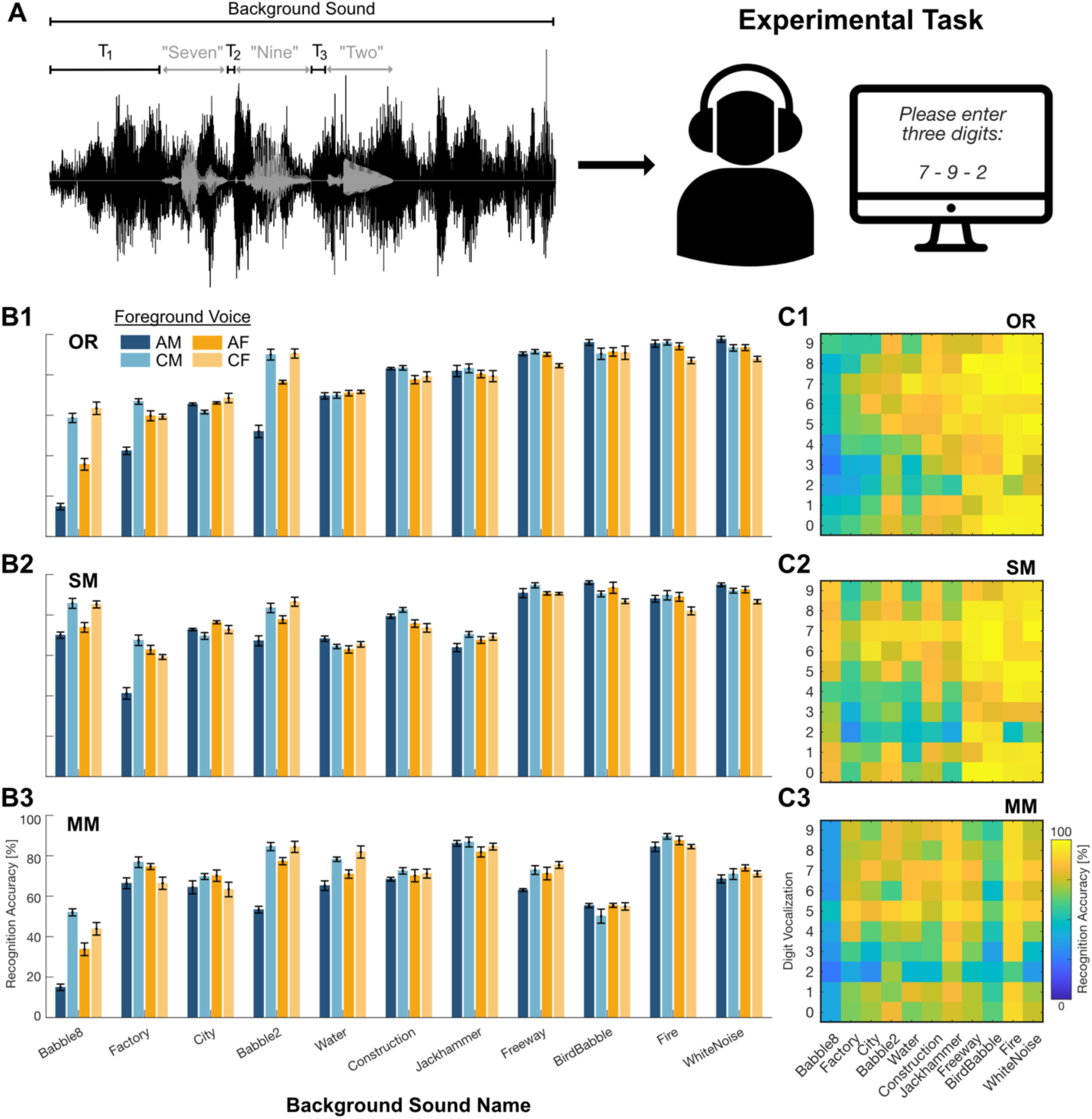
A) Participants listened to speech in noise sounds over headphones and were required to identify the digits in the sound mixture via keyboard (e.g., 7-9-2). B1-B3) Digit recognition accuracy for each voice identity (adult/child, male/female), background and perturbation (OR, SM, MM) averaged across participants (n=10). Error bars indicate sem across participants. C1-C3) Single-digit recognition accuracy for individual digits, backgrounds and perturbations (OR, SM, MM) – averaged across participants.

Accuracy also varied when the target digit was changed. For instance, with the original backgrounds (Fig. 2C1), the accuracy for the spoken digit *two* was lower across all backgrounds in comparison to the spoken digit *seven*. Certain digits, such as *seven*, *eight* and *six* had higher accuracy in backgrounds such as babble-8, factory, city, and babble-2, which were difficult backgrounds overall, while other digits *four, three* and *two* have lower accuracy. Beyond these broad trends, some digit-background combinations appear to be unexpectedly good or unexpectedly bad. For instance, *six*+*city* and *seven*+*water* have higher accuracy than expected from their marginal effects. In contrast, *three*+*babble-8* and *two*+*fire* have lower accuracy than expected. Furthermore, these trends appear to depend on both the background spectrum and modulation statistics. For instance, when the modulations are removed from the jackhammer sound (SM), digits *zero*, *three*, and *four* show lower accuracy compared to OR, whereas *seven* maintains a similar accuracy compared to OR (Fig. 2C2).

The effects of background spectrum and modulation cues can be summarized by looking at the difference in accuracy between the perturbed and original backgrounds (Fig. 3A and B). Here we compute a modulation (OR-SM) and a spectrum effect (OR-MM). As in previous work ^2^, we find that the babble-8 background has a large modulation effect where the modulations in the original sound lead to reduced recognition. Additionally, adult male and female voices in babble-8 have this same modulation masking effect, but children’s voices appear to benefit from the spectrum of the original background sound (Fig. 3A, red). In comparison, the modulations of the original jackhammer background appear to improve recognition (positive modulation effect), with minimal differences between the different voice groups (Fig. 3A, medium blue). Similar trends appear when looking at individual words, where there is large variability across words in the babble-8 condition (Fig. 3B, red), but restricted variability in the construction sound (Fig. 3B, light gray). Overall, the variation in recognition accuracy across words, speakers, and background presents a complex picture where many different acoustic factors impact recognition. However, there is substantial consistency across normal hearing participants (pair-wise correlations across all voice groups and backgrounds, r=0.92±0.002; pair-wise correlation across all digits and backgrounds, r = 0.85±0.004; mean±sem).

**Figure 3:**
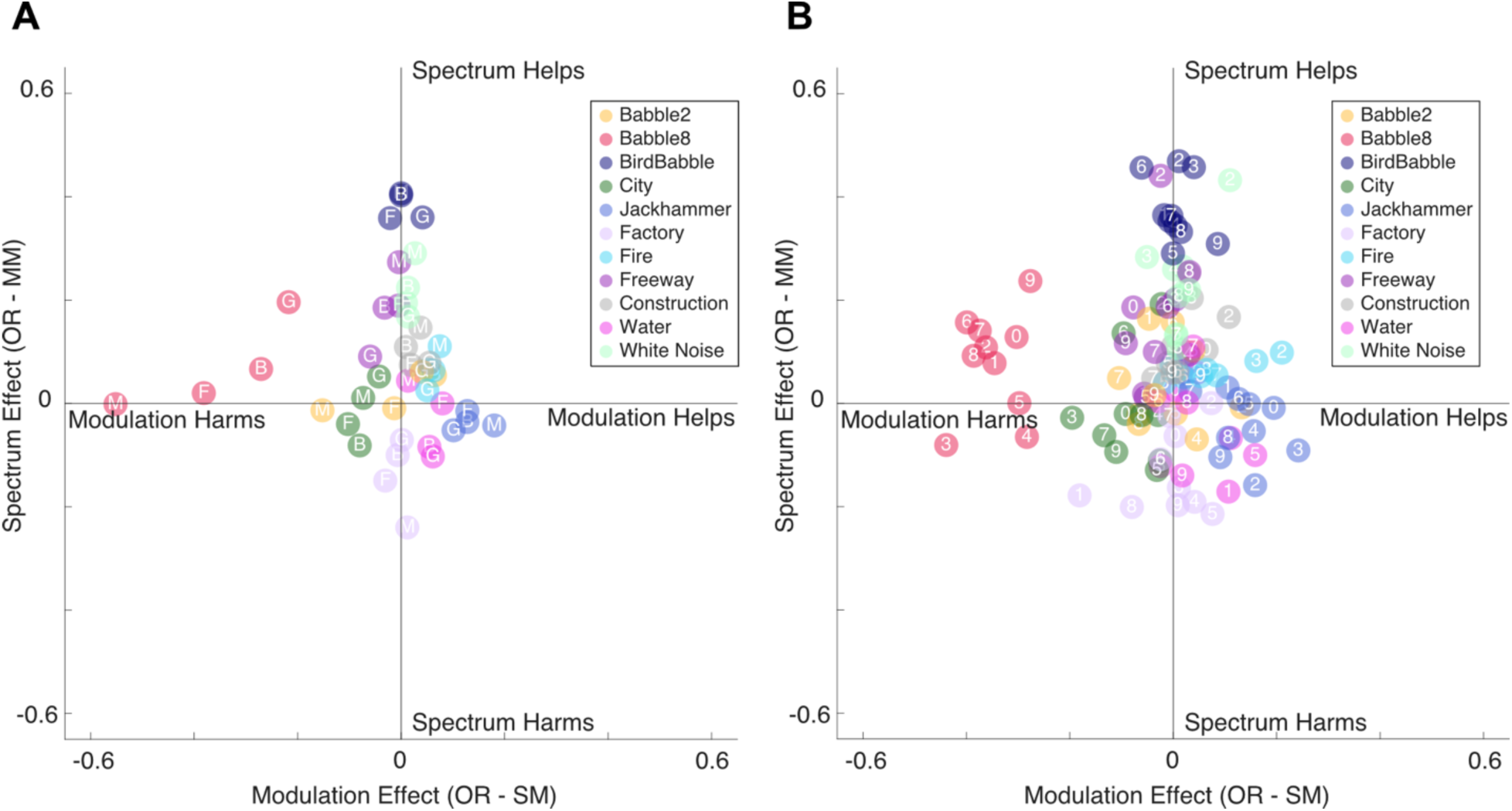
Spectrum- and modulation-driven accuracy effects. To quantify the contributions of spectrum and modulation statistics on accuracy we compared performance on OR trials to trials where the background was perturbed. The difference OR–SM indicates the change in accuracy associated with the modulations in the original background sound (Modulation Effect). The difference OR–MM indicates the change in accuracy associated with the spectrum of the original sound (Spectrum Effect). A) Modulation vs spectrum effects for each voice Adult Male (M), Adult Female (F), Child Male (B; Boy), and Child Female (G; Girl), shown for every background. B) Modulation vs spectrum effect for each foreground digit (*zero* through *nine*) for each background. Results are averages across n=10 participants.

### Recognition abilities associated with articulatory features

To better understand what foreground features might lead to accuracy differences, we extracted four articulatory features from each spoken digit: the fundamental frequency (F0), first formant (F1), second formant (F2), and harmonicity index (see Methods). Combining data across digits, we found that AM voices had lower fundamental frequency (116.8±18.4 Hz, mean±std; Fig. 4A), compared to the CM and CF groups (CM, 236.7±32.9 Hz; CF, 233.4±24.3 Hz, mean±std, mean±std ) and the AF group (201.0±27.5, mean±std). These distributions are consistent with differences in larynx size between voice groups ^30^. Distributions for F1 and F2 (Fig. 4A) are closer across voice groups but maintain a similar organization with AM voices being the lowest frequency (473.1/1469.3 Hz, mean), followed by AF voices (532.4/1689.4 Hz, mean), and then the CM (585.8/1845.6 Hz, mean) and CF (621.0/1857.5 Hz, mean) voices, as expected from the vocal tract anatomy and length ^31^. Harmonicity measures follow a similar trend with AM voices having the lowest harmonicity and with AF, CM, and CF voices having substantial overlap in harmonicity. Note that some of the overlap in these distributions, particularly the F1 and F2 distributions, is likely driven by the fact that here we are combining data from all 10 target words, which differ in their vowels.

**Figure 4:**
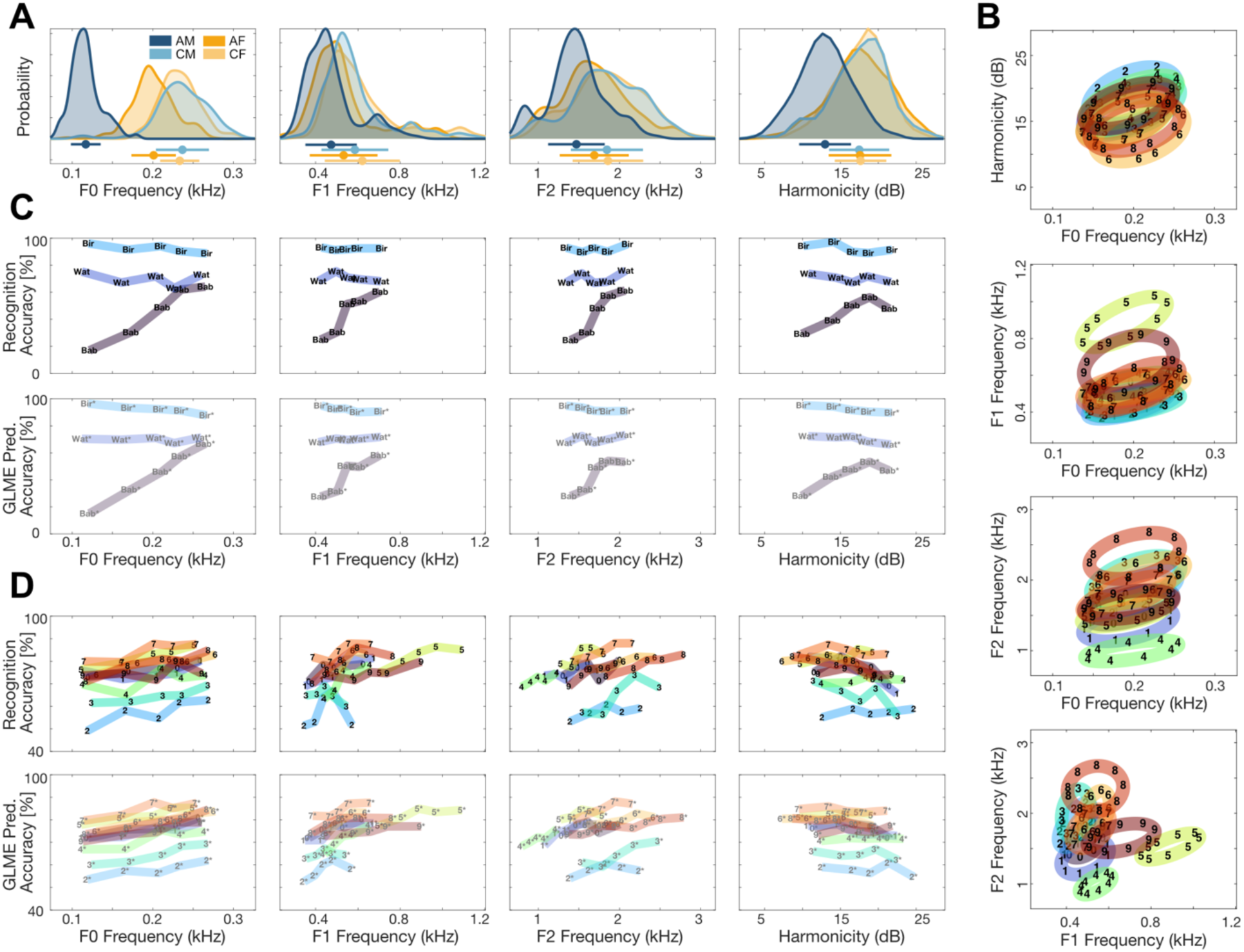
Contributions of voice features. For each foreground digit we extracted four voice features: fundamental frequency (F0), first formant frequency (F1), second formant frequency (F2) and harmonicity (HM). A) Distributions of F0, F1, F2, and HM according to voice group (adult/child and male/female). B) Bivariate Gaussian distributions fit to F0, F1, F2 and HM pairs to visualize differences across target digits. Ellipses denote 1 standard deviation and are centered on the mean. C) Participant digit recognition accuracy (top) varies across backgrounds and its associated voice features (top). Trends denote participant averages for trials split into 20% quantile bins for F0, F1, F2 and HM and for three different backgrounds: Bird Babble OR (Bir, cyan,), Water OR (Wat, blue) and Babble 8 OR (Bab, indigo). Additionally, we fit a mixed-effects model (GLME; see Methods) with fixed background and F0 effects, but without other voice features. Predictions are shown at the bottom. D) Participant accuracy (top) and GLME predictions (bottom) for the different target digits (0-9). Data is again split into 20% quantile bins for F0, F1, F2 and HM, and averaged across participants.

Next, we characterized the variability and covariability in vocal features across digits, combining data across the different voice populations. Here, average F0 does not vary substantially between digits (Fig. 4B, top) but has substantial variability across speakers – with the 20th percentile having an F0 of 110Hz and the 80th percentile 260 Hz (Fig. 4A). Harmonicity, on the other hand, does vary somewhat between digits with *two* having the highest harmonicity and *six* having the lowest (Fig. 4B, top). Additionally, we find that harmonicity appears to be positively correlated with F0 (r=0.41). F1 and F2 show substantial variation across digits, with each digit having different formant frequencies, largely driven by differences in vowels. For instance, the spoken digit *five* has a center F1 frequency of ∼890 Hz, while the spoken digit *three* appears to have a center F1 frequency of ∼440 Hz. Similarly, there appears to be a separation in F2 for different digits, where *eight* has a high center F2 at ∼2300 Hz, where *four* by contrast has a center F2 lower at ∼1000 Hz. Both F1 and F2 also have slight positive correlations with F0 (r=0.29; r=0.36), consistent with the group ordering. Comparing F1 and F2 together (Fig 4B, bottom), shows a nearly non-overlapping digit representation. Here, the spoken digit *eight* is spatially located at a center F1 ∼540 Hz and F2 ∼2300 Hz, whereas *five* is spatially located at a center F1 ∼890 Hz and F2 ∼1500 Hz. Speaker variability appears to be driven by F0, while the variability in different words comes from F1 and F2.

We then sought to explore how recognition accuracy varied as a function of the fundamental and formant frequencies and harmonicity for different backgrounds (Fig. 4C) and digits (Fig. 4D). Here, the association between speech articulatory features and accuracy depends on the background. For instance, in the presence of babble-8 noise, accuracy increases with F0 from 17% for the lowest F0 quantile to 64% for the highest F0 quantile (Fig. 4C, top, left). This is consistent with the result above, where CM/CF foregrounds had higher accuracy in the babble-8 condition compared to AM/AF. The babble-8 background is comprised of multiple adult speakers (see supplemental audio), suggesting that lower fundamental frequency voices will overlap more with the background. In contrast, in the presence of bird babble noise, accuracy slightly decreases as a function of F0 from 96% accuracy for the lowest F0 quantile to about 89% for the highest quantile. Since the frequency content of the bird chorus is higher than babble (90% percent of power within ∼0.45-3.6 vs ∼0.06-1.0 kHz ), masking may occur more for higher F0 voices. Lastly, in the presence of the water background, there is little to no change in accuracy across F0. Although these three backgrounds are only a subset of the data, they illustrate how F0 can interact with the background to determine recognition accuracy, and we find similar interactions for F1, F2, and harmonicity (Fig 4C, top).

In contrast, we find that the foreground features, while they affect accuracy, generally do not interact strongly with the digit identity (Fig 4D). Accuracy slightly increased for all digits as F0 increases (Fig. 4D, left). Accuracy also increases with increasing F1 and F2 but does not change substantially with increasing harmonicity. Although the digits differ in their F1, F2, and harmonicity distributions, and each digit has a different baseline accuracy, the effect of the different vocal features on accuracy is largely consistent across digits. The digit *two* had the lowest average recognition accuracy, while *seven* has the highest accuracy, but accuracy for both digits increases as a function of F0, F1, and F2 in a similar way. There is minimal interaction between vocal features and digit identity when describing recognition accuracy.

To quantify the magnitude of the foreground feature effects we developed a Generalized Linear Mixed Effects Model (GLME, See Methods) and identified the best fixed-effects structure using model comparison for each participant. Single-trial accuracy data was best explained by a model with fixed effects for the background category, digit category, digit position (first, second, or third), with an effect of F0, and with background*F0 interactions, but no effects of F1, F2, or harmonicity. The predictions of this GLME (Fig. 4C, bottom) are shown for accuracy across F0, F1, F2, and harmonicity. At least in this data, the associations between foreground features and accuracy across backgrounds (Fig 4C) and across digits (Fig 4D) appear to be largely explained by F0 alone. There are also substantial differences across backgrounds (F(10,98923)=132.48, p<0.001, 0.10-686.3, odds ratio range referenced to babble-2) and digits (F(9,98923)=357.82, p= p<0.001, 0.33-1.91, odds ratio range referenced to *zero*). We also find that the order of the digits plays a role (F(2, 98923)=70.94, p<0.001), with recognition accuracy being higher when the digit is in the 2nd (odds ratio 1.10) or 3rd position (odds ratio 1.28). The main effect of F0 was substantial (F(1, 98923)=229.52, p<0.001) with an odds ratio of 1.02/Hz, and a statistically significant interaction with the background (F=66.10, p<0.001).

### Subband modulations preserve articulatory feature representations

Next, we test whether spectrotemporal summary statistics can account for speech articulatory features among a wide range of male and female talkers of multiple ages. Here, we use a biologically motivated, hierarchical model of the auditory midbrain containing spectrotemporal receptive fields that allow for measuring the modulation statistics for each spoken digit (Fig. 5A, See Methods). The model output statistic, the *midbrain modulation power spectrum* (mMPS), accounts for the sound’s power distribution in three acoustic dimensions: frequency, temporal and spectral modulation. To visualize how well articulatory features are preserved in this statistic, we applied t-distributed stochastic neighbor embedding (t-SNE) to the mMPS to represent each digit in a low dimensional 2D space.

**Figure 5:**
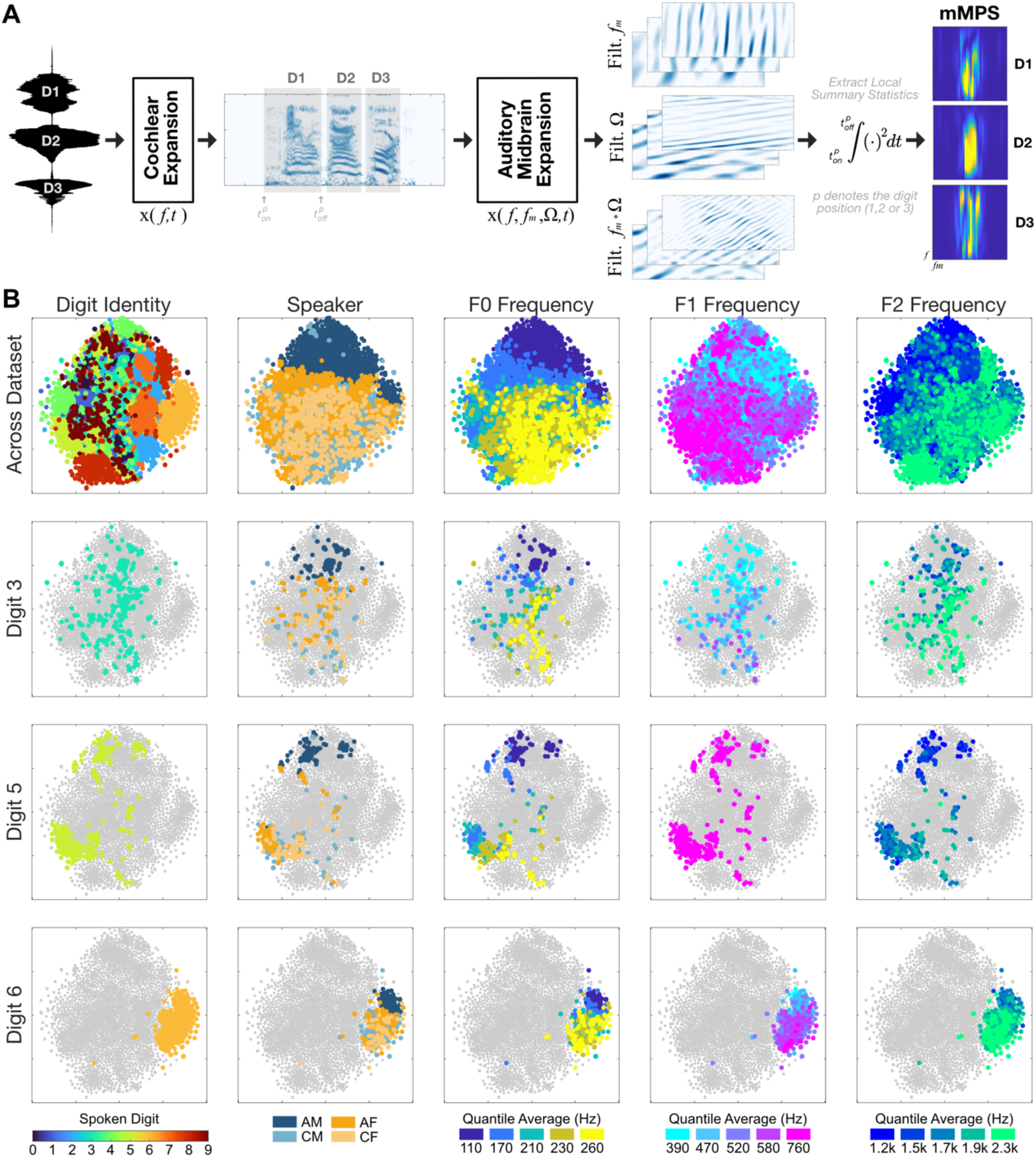
Auditory midbrain model statistics capture foreground digit information and voice features. A) Speech sounds were characterized with a two-stage hierarchical auditory model composed of 1) logarithmically spaced frequency-tuned filters that mimic the auditory periphery and 2) logarithmic spectral and temporal modulation filters that mimic the decomposition of the auditory midbrain. The resulting auditory model outputs, subbands, contain the time-varying envelopes for each frequency, temporal and spectral modulation channel. By computing the time averaged power for each subband we estimate the midbrain modulation power spectrum (mMPS) summary statistic of each sound (right), which represents the sound power density along frequency, temporal and spectral modulation. B) T-distributed stochastic neighbor embedding (t-SNE) was used to perform dimensionality reduction on the auditory representation by projecting the mMPS of each sound onto a 2-D spatial mapping that groups sounds according to their mMPS similarity. We then visualize mMPS features as a function of word and articulatory features (e.g., digit, speaker, F0, etc.; left to right). The overall organization for digit, voice group, F0, F1, and F2 are shown on the top row. Bottom rows highlight foregrounds from individual digits (3, 5 and 6) with all other digits in gray. The dot positions in all plots are identical.

In the embeddings, we find that different digits and voice groups are well differentiated (Fig 5B, left) and there is substantial variability in the clustering across digits. For instance, the spoken digit *three* spans a broad range of the embedding space, while *six* is localized to a small part of the space. The mMPS features are also differentiated by voice group in the embedding at the level of the whole dataset and within individual digits. The embeddings of the different voice groups are mirrored by the patterns across F0 and, to a lesser extent, F1 and F2 (Fig 5B, right). In the mMPS space, articulatory features vary systematically across the different word and voice categories. Since the mMPS preserves differences in both articulatory features and digits, it may be able to account for the behavioral trends described above.

### Subband modulation masking explains variability in recognition accuracy across words and voices

To determine whether the mMPS can account for both foreground- and background-driven changes in recognition accuracy, we next apply a previously developed, generalized perceptual regression (GPR) approach ^2^ (Fig. 6A; see Methods). Here the background and foreground mMPS (Fig. 6A, left) and their interaction (point-wise products) are dimensionality reduced using principal components analysis (PCA). The resulting neural model embeddings (Fig. 6A, middle), here with 15 principal components each, are then used as input features to a logistic regression model that predicts whether the participant identifies the digit correctly (see Methods).

**Figure 6:**
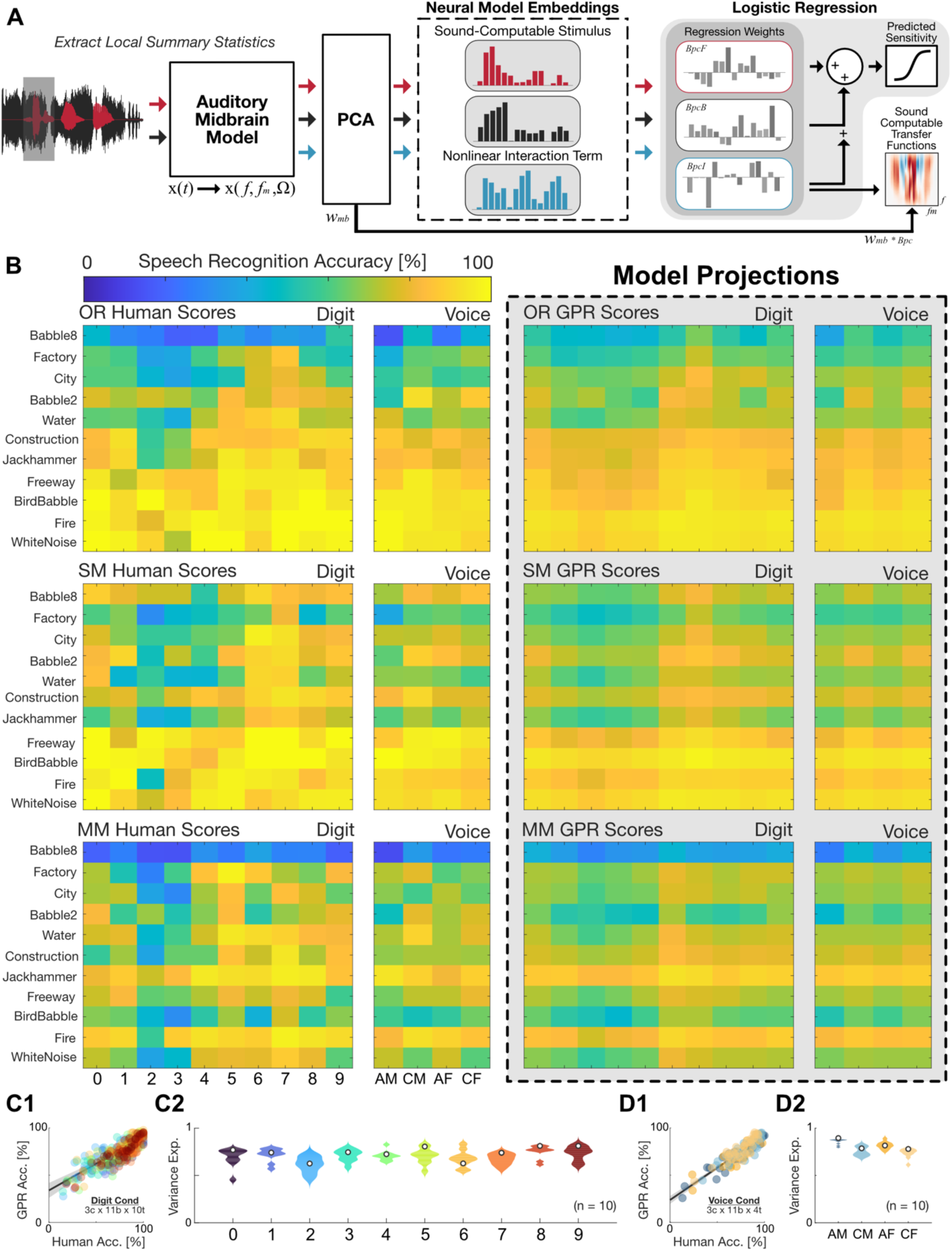
Generalized perceptual regression (GPR) applied to auditory midbrain model statistics predicts speech recognition in noise behavior. A) To predict single-trial digit recognition outcomes, we use the auditory midbrain decomposition to first extract local summary statistics (mMPS) from each foreground digit (red) and its overlapping background (black), and multiple the two mMPS to create an interaction term (cyan). Each term is then dimensionality reduced using principal component analysis (PCA). Low-dimensional features, along with the serial position of the digit are then used as input predictors to a logistic regression model that is optimized using behavioral measurements to predict the likelihood of correct decisions. The principal components and model weights can be used to derive perceptual transfer functions that indicate which specific mMPS features improve or reduce recognition accuracy. B) Average single-digit recognition accuracy scores for an individual participant (participant 9) are shown for each background and perturbation (OR, SM, MM, top to bottom) for both the digit (left) and voice projections (right). Right panels show cross-validated GPR model predictions for the individual participant (model constrained to 15 PCs for each input). C1) Observed vs predicted accuracy for each digit and background for the example participant. C2) Violin plots (right) show variance explained across participants for each digit, white dot represents the example participant. D1) Observed vs predicted accuracy for each voice group and background for the example participant. D2) Violin plots (right) show variance explained across participants in each voice group, white dot represents the example participant.

We evaluated cross-validated predictions for individual participants’ recognition accuracy across a digit projection (DP) where the data and predictions are averaged across speakers and a voice projection (VP) where the data and predictions are averaged across digits (Fig 6B, shown for one participant). GPR accurately replicates the overall patterns in recognition accuracy across different voice groups, spoken digits, and backgrounds. For instance, in the case of the original sounds (Fig 6B, top left), the GPR scores replicate the pattern of low accuracy for babble-8, factory, city and water for most digits, as well as trends across different voices and backgrounds where AM/AF have a lower accuracy than CM/CF for babble-8, factory, city, and water backgrounds. The model also captures accuracy differences in SM (Fig 6B, middle) and MM (Fig 6B, bottom) backgrounds, which are driven by changes in the background spectrum or modulation content. Together, these predictions suggest that spectrotemporal modulations, and the interaction between foreground and background mMPS features, are sufficient to reconstruct human perceptual performance in the task. For this individual participant, the GPR explains 73% of the perceptual variance across digits (Fig. 6 C1 and C2, white dots) and 78% of the perceptual variance across voice groups (Fig. 6 D1 and D2, white dots).

Across participants, the variance explained by the model for the digit projection ranged between 66.0-73.2% (69.4± 0.8%, mean±sem; Fig. 6C2), with the digit *two* having the lowest explained variance for the digit projection (62.5±0.1%; mean±sem) and *eight* having the highest prediction quality (75.4±0.1%). For the voice projection the model explained 75.6 – 83.3% of the variance across participants (80.2 ± 0.9%, mean±sem , Fig. 6D2), with the lowest prediction quality for CF voices (73.6±0.1%) and the highest for AM voices (85.8±0.1%).

GPR also provides an interpretable framework that allows visualizing the specific frequency and modulation features, in both the foreground and background, that influence accuracy. Here, the participant-fitted model regression weights are interpreted as *perceptual transfer functions* (pTF; see Methods) that indicate each listener’s sensitivity to foreground and background modulation features (Fig. 6A, bottom right). The foreground and background pTF are shown for the 0 cycle/oct spectral modulation, which dominates the perceptual sensitivity^2^. The pTFs of different participants and the population (Fig. 7A) share similar structure (inter-participant correlation: foreground=0.92±0.01, background=0.97±0.01; mean±sem) and cover a broad range of frequencies. However, the pTF extent in the temporal modulation dimension differed between the foreground and background. For the foreground, pTFs were restricted to below ∼8 Hz, consistent with prior studies showing that slow temporal modulations are critical for speech recognition ^23^. By comparison, the background temporal modulations have a broader influence on accuracy, since visible components in the background pTF can exceed 64 Hz (Fig. 7A).

**Figure 7:**
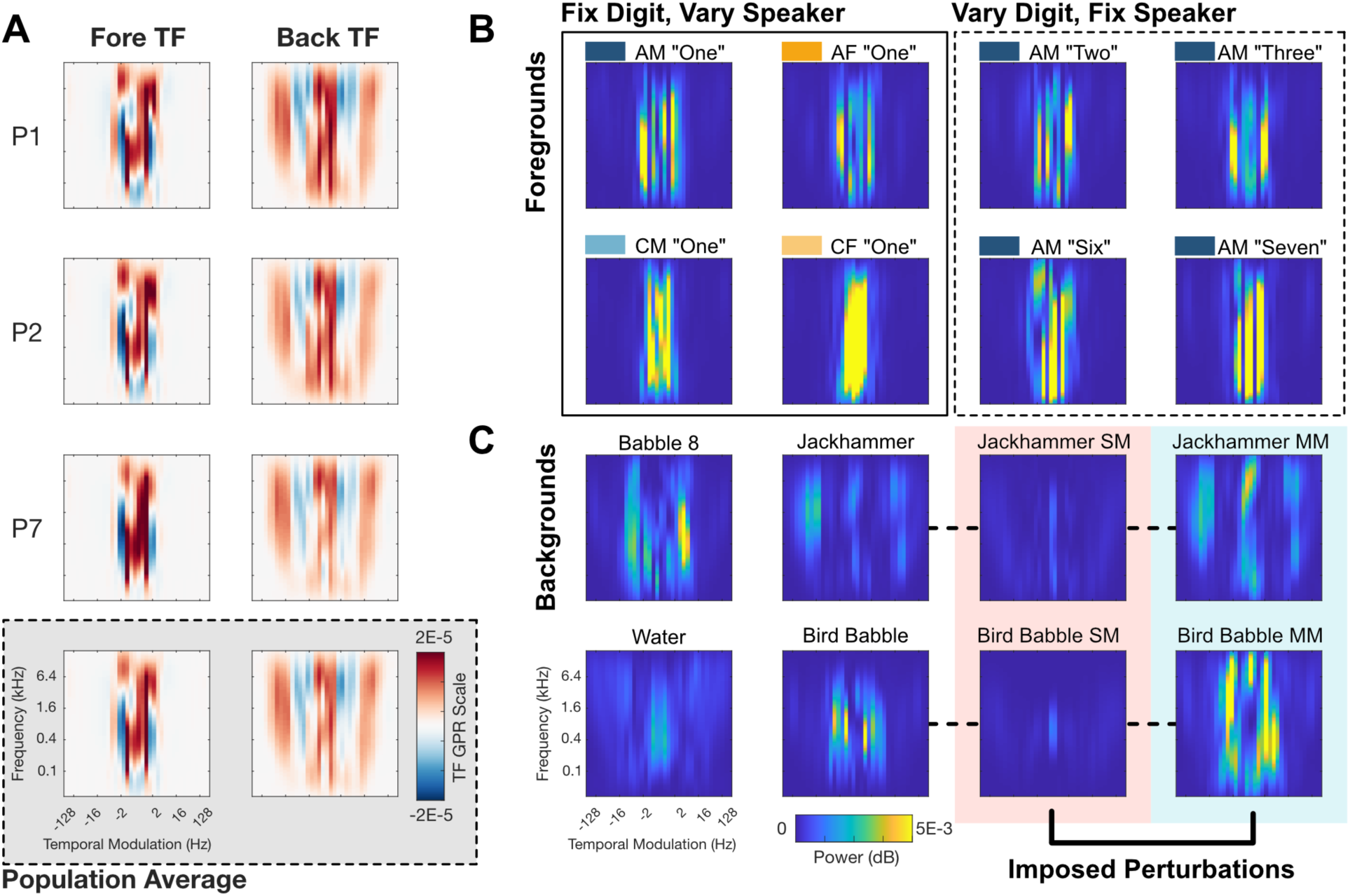
Generalized perceptual regression (GPR) weights and model comparison. A) Using the principal components and GPR model weights, we calculate a *perceptual transfer function* that can be used to interpret the contribution of individual mMPS components (see Methods). Transfer functions are shown at ripple density 0 cycles/oct for three participants (top) alongside the population average (bottom) for both the foreground (left) and background (right) features. Positive weights (red) indicate features that improve accuracy while negative weights (blue) indicate features that decrease accuracy. Weights for other spectral modulation frequencies and the foreground x background interaction term are omitted for clarity. B) Example foreground stimuli shown in the same mMPS space (ripple density 0 cycles/oct). C) Example background stimuli shown in the same mMPS space for different backgrounds (left) and their corresponding perturbations (right).

The foreground and background pTFs also exhibit frequency and modulation specific excitatory (increased accuracy) and suppressive (decreased accuracy) domains that provide a direct description of perceptual sensitivity. Predicted accuracy is derived as a dot product between the foreground mMPS (Fig. 7B) and its pTF (Fig. 7A, left) and the background mMPS (Fig. 7C) with its pTF (Fig. 7A, right), followed by a logistic nonlinearity. Background-driven accuracy differences can be explained by the combination of the stimulus mMPS and the participant’s pTF. For instance, compared to the OR and SM conditions, the MM bird-babble (Fig 7C, bottom row) has very high power directly overlapping the background pTF inhibitory domain (∼2-8 Hz; Fig. 7A, right), producing a lower predicted accuracy than the SM and OR conditions. The OR and MM jackhammer background (Fig. 7C, top row), in contrast, contain high power overlapping the excitatory pTF domains (∼16-64 Hz temporal modulation; Fig. 7A, right), producing a high accuracy. For this same background, the SM manipulation substantially reduces the power that overlaps the pTF excitatory domains, thus reducing predicted accuracy. Across speakers or digits, the foreground pTF is similarly combined with the foreground mMPS to account for the measured accuracy. For a fixed digit, the mMPS can vary substantially across speakers (Fig. 7B, left) and, for a fixed speaker the mMPS shows substantial word-related variability (Fig. 7B, right). While these sounds represent only a small fraction of sounds used, this variability in the mMPS across words and speakers naturally leads to many possible interactions with the excitatory and suppressive domains in the foreground pTF (Fig. 7A, left), reducing or increasing the predicted recognition accuracy.

### Contributions of spectrum & modulation, foreground & background, and glimpsing

We also consider several reduced GPR models to determine which background and foreground statistics contribute to speech recognition accuracy in noise. The full GPR model takes the background and foreground mMPS and its nonlinear interaction as predictor variables (mMPS*_b_*+mMPS*_f_* + MPS*_b_* × MPS*_f_*). To determine how each of these terms contributes to the prediction quality we selectively remove different components from the model (Fig. 8A). Cross-validated prediction quality for the voice and digit projections as well as the single-trial predictions follows a rank order where the full model has highest accuracy, followed by the mMPS*_b_*+mMPS*_f_* model, mMPS*_b_* model, and finally the mMPS*_f_* model configuration. Including both the foreground and background mMPS accounts for most of the model predicted variance, although the background only model is far more predictive than the foreground only model. Thus, the background statistics provide the largest contribution to perceptual sensitivity, while the foreground statistics have a smaller, but reliable, influence that allows for word- and speaker-related refinements to prediction accuracy.

**Figure 8:**
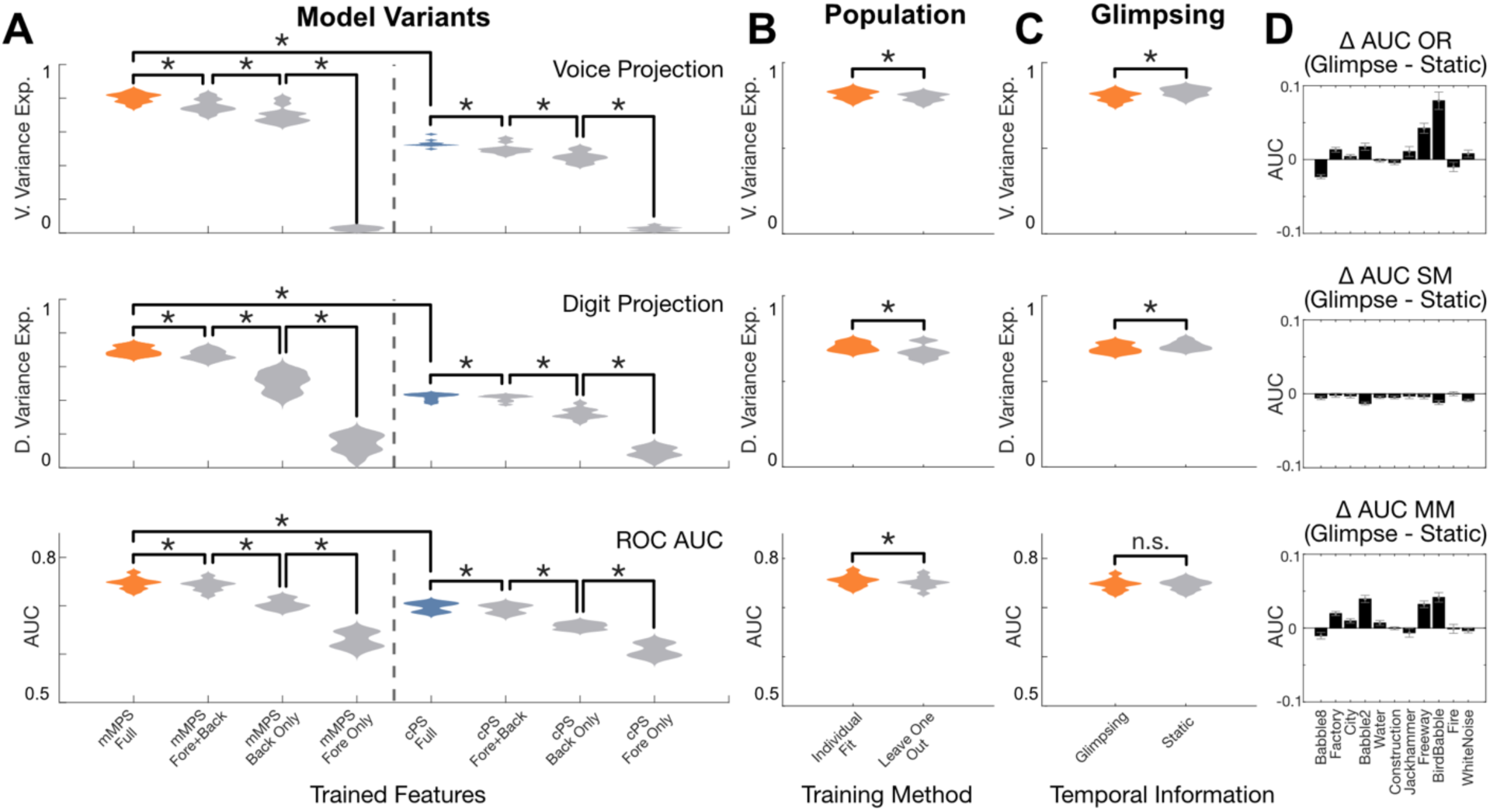
GPR model goodness-of-fit along with comparisons for reduced models. A) Violin plots indicate results across participants for variance explained in average digit recognition accuracy across voice group, background, and perturbations (voice projection, top), across digit, background, and perturbations (digit projection, middle), and single digit accuracy area-under-the-curve (AUC). Reduced models vary in either 1) task features (Foreground, Background, Interaction) and/or 2) auditory features (midbrain features, mMPS, or cochlear features, cPS). B) Violin plots show goodness-of-fit results for the full (mMPS interaction) model across participants for individually fit and population level models (leave-one-participant-out cross-validation). C) Violin plots show results for a model that accounts for local summary statistics (glimpsing), and a model that uses average summary statistics (static). D) Change in AUC (glimpsing – static) for each background and perturbation (OR, top; SM, middle; MM, bottom) combination.

In addition to characterizing sounds and perception with the mMPS we also consider the cochlear power spectrum (cPS) as the input predictor variable to the GPR^2^. The cPS statistic accounts for power density of the sound as derived through our cochlear model, such that, it contains frequency related information but lacks information about the modulation statistics. The cPS predictions across the full and reduced models (Fig. 8A, right) follows a similar trend for the mMPS model (Fig. 8A, left; VP: 53.1±0.7%, DP: 42.2 ± 0.6%, AUC: 69.8±0.3). However, the cPS model has significantly lower predictive power than that of the mMPS (VP Fisher z-transform paired t-test, t(9) = 21.1, p < 10^-8^; DP Fisher z-transform paired t-test, t(9) = 25.85, p < 10^-9^; AUC paired t-test, t(9) = 29.60, p < 10^-9^), consistent with our prior findings^2^. Thus, although the frequency content of each sound contributes, the modulation statistics are critical to explain perception.

We also examine generalization across participants by testing whether a population average model can account for perceptual sensitivity from held-out participants. Here we find that the individual participant models are significantly better than the population average model predictions (VP: t(9) = 7.4, p < 10^-4^; DP: t(9) = 3.78, p < 10^-2^; AUC t(9) = 10.18, p < 10^-5^). However, the individualized models provide relatively small improvements in prediction quality (Fig. 8B, VP: 81.4±0.8% vs. 79.7±0.7%; DP: 70.2±0.8% vs. 67.5±1.2%; AUC: 75.5±0.4 vs. 75.0±0.4), suggesting that recognition patterns are largely consistent across this sample of typically hearing participants.

Finally, we tested whether glimpsing^14,15^ contributed to the perceptual trends and model predictions. By design, the background mMPS used in the full model is computed for the sound segment directly overlapping each digit, so that it accounts for momentary changes in the background sound power and has the potential to account for glimpsing effects. Here, rather than using these time-localized mMPS, we consider a time-averaged mMPS for the background as input to the GPR model. This “static” version of the model does not account for momentary changes in background power and, thus, it cannot account for glimpsing effects. Here we find that the average prediction quality for the voice and digit projections and single trials of the static model is comparable to that of the full glimpsing model (Fig. 8C; VP: 82.8±0.8% vs. 80.2±0.9%, DP: 70.7±0.8% vs. 69.4±0.8%, AUC: 76.6±1.0 vs. 74.7±0.4). However, notable differences in the prediction quality can be observed for the predictions within individual backgrounds (Fig. 8D).

Glimpsing effects, where there is a difference between the original and static models, appear to be exclusive to nonstationary background sounds such as the freeway noise (sparsely passing cars), two-speaker babble, and bird babble (a chorus of 2-3 geese). For each of these sounds, there is an increase in the prediction quality for the glimpsing model compared to the static version, and this effect is particularly noticeable for the MM condition (bottom). In contrast, the glimpsing and static model perform near-equally well for the SM sounds (middle). This is expected since the SM sounds have whitened modulations and are stationary by design.

## Discussion

The findings demonstrate that the spectrum and modulation statistics of speech interact with those of a competing background sound to influence word recognition accuracy. Beyond the interference effects driven by the background sound statistics^2^, word-level and speaker-dependent articulatory features, including the voice fundamental frequency and formants, have a more nuanced influence on accuracy. These articulatory features vary systematically between words and speaker groups, and we demonstrate that low-dimensional auditory model embeddings capture word and voice related information. More broadly, a low-dimensional regression model that uses auditory modulation statistics produces accurate behavioral predictions across 9,900 sounds encompassing many talkers, backgrounds, and words on a trial-by-trial basis. Altogether, these results demonstrate how using modulation statistics in a generalized perceptual regression (GPR) model framework can provide a biologically interpretable account of speech recognition behavior in highly variable and realistic listening scenarios.

How humans solve the *cocktail party problem* ^1^ remains an important challenge in auditory neuroscience, since everyday hearing invariably occurs in cluttered auditory scenes and such scenarios severely impair abilities for those with hearing loss ^32,33^. Prior studies have focused mostly on perception of controlled speech stimuli and simplified backgrounds that lack diverse statistical structure of real-world backgrounds ^22,34,35^. Although these studies have highlighted how specific foreground or background features can impact speech recognition, limited stimulus variability may make it difficult to identify the broader representations that underlie hearing any speech in any noise.

Our findings here suggest that spectrum-based interference, or energetic masking ^16^, alone is insufficient to fully explain voice, word, and background related effects and that spectrotemporal modulations may allow a generalizable explanation for speech perception in its full variability. Of all the factors tested, the background sound has the strongest impact on accuracy, yet word-level and speaker-level statistics interact with the background to modulate perceptual accuracy. Additionally, although speech articulatory features, such as the fundamental and formant frequencies, can account for word and voice driven recognition effects (Fig. 2), here we find that these features need not be explicitly computed to explain behavioral findings. Instead, a biologically plausible representation involving frequency and modulation selective filters that mirror the representation in subcortical regions can explain behavioral findings across diverse backgrounds, talkers, and words on a single trial basis. This is consistent with a computational framework whereby interference occurs within specific subbands, with spectrotemporal filters that are jointly selective for frequency and spectrotemporal modulations. The auditory midbrain is likely a key brain region where such a decomposition and any interference would occur, since it is the first stage in the auditory system exhibiting a preponderance of modulation tuning ^36^, covering an extensive range of spectrotemporal modulations (to ∼500 Hz) ^36,37^. This is in contrast to cortical regions, which have a more restricted range of modulation selectivity (mostly < 20 Hz) ^36,38,39^ and would be less likely to account for modulation-driven behavioral effects shown here.

Our approach is related to several existing measures of speech intelligibility^40,41^. Although these previous measures have focused on speech quality ratings after distortion or compression, they often use similar statistical features to those used here. There are a wide variety of sound representations, and modulations are likely to play a key role in many tasks^23,42–45^. Our results also provide an alternative and complementary modeling framework to task optimized deep neural network models ^46–48^. Such models can replicate some aspects of human behavior but often have tens of layers and thousands to millions of parameters. The parameters of neural network models can also be difficult to interpret and attempts to probe such networks have revealed that these models can differ substantially from human perception. For instance, task-optimized neural networks seem to use different invariances for sound recognition compared to humans^49^, and the modulation statistics influencing machine perception are not well matched to human speech-in-noise behavior^2^. Unlike neural networks, which are optimized to perform a particular task (e.g., predict words), GPR is optimized to predict a listener’s behavioral sensitivity (correct or incorrect judgments). In this manner, GPR takes the sound representation and predicts behavioral outcomes, requiring few dimensions (45 here) and producing linear perceptual transfer functions for each listener that directly indicate which modulation statistics impact perception. Since the GPR framework produces a listener-specific model of hearing and identifies the statistical features underlying individual behavior, it could potentially be applied to compare perception across people, animals, or machine systems and could also be used as a diagnostic for hearing disorders^43^ or speech perception difficulties.

Although our results here provide a new, generalizable model of speech in noise perception, there are several limitations for our approach and important open questions. First, our findings are based on semi-realistic listening conditions using natural background sounds and speech consisting of multiple words, talkers, and voice categories. While these experimental sounds are acoustically diverse, natural auditory scenes also contain variability in reverberation and spatial cues that distort spectrotemporal content and alter speech perception ^18–21^. It is important to note that here we also presented sounds at a fixed SNR and intensity. Testing perception under more variable conditions could potentially alter the model performance. The model itself is also based on trial-wise prediction. Using a convolutional GPR^43^ , with unknown word onsets and offsets, may better account for the temporal dynamics of continuous speech perception, such as glimpsing and co-articulation. Finally, neural and behavioral measurements alongside GPR may provide a more thorough test of how spectrotemporal modulation cues are represented for sound mixtures.

## Materials and Methods

### Human Psychoacoustics

Experiments were approved according to the University of Connecticut Institutional Review Board (IRB). We recruited female (n=7) and male (n=5) native English speakers, ages 21-47 who had normal hearing sensitivity (>20 dB HL for left and right ears, 0.25, 0.5, 1, 2, 4, 8 kHz, tested with a GSI Audiostar Pro audiometer [Grason-Stadler], with a Radioear 3045 supra-aural headset). Participant responses were recorded with anonymized participant IDs for confidentiality. Of the (n=12) enrolled participants, (n=2) did not complete the study and were thus excluded from the data analysis. The experiments were conducted at the University of Connecticut Storrs Campus and in a sound shield room (Industrial Acoustics Co.). Sounds were delivered through a Fireface UC digital audio interface (RME Audio Inc.) amplified with a BHA-1 headphone amplifier (Bryston Ltd.) and sent to HD 820 headphones (Sennheiser Co.). All sounds were calibrated to 65 dB SPL. Sound delivery and the acquisition of behavioral responses via keyboard entry were controlled using custom software using Psychtoolbox ^50^ in Matlab (Mathworks Inc.) on a Mac Mini M1 computer (Apple, Inc.).

Participants were presented with sound mixtures consisting of foreground speech and background environmental noises to assess how both foreground and background features impact recognition performance in natural auditory scenes. Each speech sequence (Fig. 1C) consisted of a randomly chosen triplet of English digits (D1-D2-D3), delivered with a random inter-digit time interval during each trial (T1-T2-T3) and with different speakers (S1, S2, and S3). During each trial the three speakers were randomly selected from one voice group population: Adult Male (AM), Child Male (CM), Adult Female (AF), or Child Female (CF) (see Fig. 1E). Speech digits were selected from the TIDIGITS corpus ^51^ which contains 326 talkers (111 AM; 114 AF; 50 CM; 51 CF). Each foreground triplet sequence was paired with a natural or perturbed background masker at an SNR of -9 dB. The instantaneous timing of each digit was used to calculate the SNR, such that the silent gaps (intervals between digits) did not contribute to the SNR measurement, which would create an artificially lower SNR. The onset and offset segments of all sounds were windowed with a 100 ms B-spline ramp to minimize transients at the beginning and end of each sound. For each sound, participants were tasked with reporting the three digits sequence that was presented.

### Background Sounds and Manipulations

Ten natural background sounds (speech babble, running water, traffic noise, jackhammer and factory sound, etc.) and white noise (control) were used as masker stimuli, similar to previous experiments^2^. For some trials, we altered the statistical structure of the background stimuli to characterize how the spectrum and modulation statistics of the backgrounds specifically influence recognition. As a reference, natural backgrounds were presented in their unaltered original (OR) state with their full acoustic structure. We also delivered spectrum matched (SM) variants which were generated by computing the Fourier transform, preserving the magnitude spectrum and randomizing the phase spectrum, and then using an inverse Fourier transform. Although SM backgrounds have power spectra that are matched to the OR, they have a whitened modulation power spectrum (Fig. 1D). We also generated modulation matched (MM) variants of each background. Here the power spectrum of the background sound is equalized to a 1/f power spectra in the Fourier domain while the phase spectrum is preserved. This manipulation alters the spectral profile of each background, while maintaining (or matching) the modulation statistics (Fig. 1D). The SM manipulation is analogous spectrum shaped noise, in which the identity of the sound is ambiguous. In contrast, MM variants maintain the original time varying fluctuations of the OR stimulus, and sound categories can still be identified. MM variants of multi-speaker babble, for instance, have identifiable speech content but with noticeable different timber compared to the OR variant.

### Experiment Block Structure

The experiment was broken up into 25 experimental blocks, each containing 25 different trials for each stimulus condition (Fig. 1E). Each participant completed two experimental blocks at a time, typically comprising 6 hours of testing split in two 3-hour sessions. In each experimental block, each of the 11 backgrounds and their 3 perturbations were tested (33 background sounds) across the 4 voice conditions shown in Fig. 1E such that there were 132 trials per block and 3,300 trials across the experiment. Each foreground stimulus was constrained to a randomly selected set of 3 speakers (S1-S2-S3) from the designated speaker population (AM, CM, AF, CF), random time delays (T1-T2-T3) and a random digit triplet (D1-D2-D3).

### Scoring Criteria, Voice and Digit Projections

Single digit recognition performance was scored according to digit identity and serial position with 9,900 single digit measurements total for each participant. On a single trial with a target foreground digit sequence of 9-8-3, for instance, a response 8-8-1 is scored as 0-1-0 (1=correct; 0=incorrect). We then averaged single digit recognition accuracy across subsets of trial conditions to identify variations in accuracy (Fig. 2). First compute a “Voice Projection” (VP) by sub-selecting trials corresponding the unique combinations of Background (11 Natural Backgrounds), Perturbation (OR, SM, MM), and Voice Identity (AM, CM, AF, CF), with averages calculated for each participant averaging over digits and repeated trials. We then calculated a “Digit Projection” (DP) where single trial responses were averaged across unique combinations of Backgrounds (11 Natural Backgrounds), Perturbations (OR, SM, MM), and Digits (0-9). Note that trials are balanced across voice-background-perturbation conditions and unbalanced (but uniformly distributed and randomized) across digit-background-perturbation conditions. That is, voice-background-perturbation conditions have identical numbers of trials, but digit conditions do not.

### Characterizing how Articulatory Parameters Contribute to Speech in Noise Accuracy

To explore how articulatory parameters of the different digits and voices contributes towards digit recognition accuracy, we also projected accuracy as a function of the fundamental frequency (F0), first formant (F1), second formant (F2), and harmonicity (HM) of the vocalization of the measurement. We used the FastTrack plugin for PRAAT ^52^ to calculate F0, F1, F2, and HM for each foreground digit. The FastTrack PRAAT plugin was run using the default parameters, and the recommended highest and lowest frequencies based on voice groups (Adult Males 4500-6500 Hz, Adult Females 5000-7000 Hz, and Children 5500-7500 Hz). The time varying outputs over the active F0 window identified by PRATT were used to estimate articulatory parameters, where F0, F1, F2 and HM are defined as the median value of PRATT’s time varying outputs across each digit.

We first explore how the articulatory features vary across voice and word identities. First, we estimated the marginal distributions (kernel density estimates) for F0, F1, F2, HM, conditioned on the speaker identity. These conditional distributions depict the variability in the dataset accounted by the articulatory parameters for different speaker groups. Next, we estimated joint conditional distributions of the articulatory parameters for each of the spoken digits. This was done by fitting multi-variate gaussian mixture distributions jointly for F0/HM, F0/F1, F0/F2 and F1/F2 combinations, for each of the spoken digits.

Next, we projected the accuracy across participants onto the F0/F1/F2/HM distribution for 1) select backgrounds and 2) different digits. First, we sub-selected participant responses from trials in all digits and voices, at each background, calculated 5 different quantiles of F0/F1/F2/HM measurements within each set to calculate accuracy at different backgrounds covarying with F0/F1/F2/HM intensity. Secondly, we sub-selected participant responses from trials in all backgrounds and voices, at a specific digit, calculated 5 different quantiles of F0/F1/F2/HM measurements within each set to calculate accuracy per digit, covarying with F0/F1/F2/HM intensity.

### Generalized Linear Mixed Effects Model

To characterize the relationships between articulatory features, background conditions, and digit recognition accuracy, we fit a generalized linear mixed effect (GLME). Using data from all participants and trials, we aimed to describe all n = 99,000 correct vs incorrect digit recognition judgements. Here we used the ‘fitglme’ function (MATLAB 2022b) with binomial observations and a logistic link function with random effects (intercepts) for each participant listener and each TIDIGITS speaker (n=326).

We compared multiple models for the fixed effect structure including categorical descriptions of the background and perturbation, as well as, categorical descriptions of the foreground, including digit category and digit position (D1-D2-D3), and the continuous articulatory features F0, F1, F2, HM. We evaluated the model goodness-of-fit using the Akaike information criterion (AIC). We hypothesized that accuracy was driven by background effects interacting with the pitch of the speaker and modulated by the identity of the target word. To parametrize the model, we ran different variants testing this theory altering the speech articulation measurements and categorical labels and selected the model with the best Akaike information criterion (AIC). This selection ensured that our model maximized predictive power, while maintained minimal complexity. The best model, among those tested, contained fixed effects for target digit and digit position and interaction terms between the background, perturbation category, and F0.

### Local Mid-Level Acoustic Feature Extraction

In addition to describing single trial digit recognition in terms of background categories and foreground articulatory features, we also sought to evaluate whether performance can be explained by the interference of more general foreground and background sound statistics. To develop a sound computable measurement of brain-inspired auditory processing, we decomposed sounds through a two-stage hierarchical model of the auditory pathway ^35^. Target vocalizations and background noise excerpts were first expanded through a cochlear model filter bank. The cochlear output is comprised of 82 overlapping frequency-selective gammatone filters (20-20k Hz, 50% overlap, equivalent rectangular bandwidths), followed by nonlinear rectification and lowpass filtering (750 Hz) designed mimic hair cell rectification and synaptic filtering in the cochlea. The output, referred to as the cochleogram, represents the transformation from the waveform *X*(*t*) to band-limited envelopes *S_c_*(*t*, *F_l_*), where *F_l_* represents the frequency of the *l*^th^ channel.

The second stage of the hierarchical model is designed to account for spectrotmeporal modulation filtering described in auditory midbrain. Here, the cochleogram *S_c_*(*t*, *F_l_*), is passed through a set of spectrotemporal modulation filters spanning 0.5-512 Hz temporal modulations (*f_m_*, in 1/2 octave steps) and 0.25-8 cycles/octave spectral modulations (Ω*_n_*, 1/2 octave steps). Thus, the resulting output tensor, *S_m_*(*t*, *F_l_*, *f_m_*, Ω*_n_*), represents individual envelopes at the output of a given, frequency, temporal modulation, and spectral modulation channel.

To estimate time-averaged local summary statistics from each sound, we first generate the time-varying midbrainogram representation (*S_m_*) for the entire 3.5 second sound in each trial. We use the onset and offset time of each digit D*_p_* where *p* denotes digit position 1, 2, or 3 to temporally segment *S_m_* in order to measure the local power of both the foreground and background:

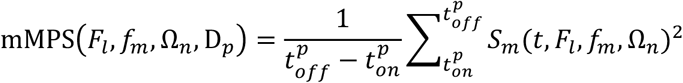

where *t_on_^p^* and *t_off_^p^* represent the onset and offset time of the *p*^th^ digit (Fig. 5A). This local summary statistic accounts for the sound power distribution across frequencies, temporal and spectral modulations as estimated using a midbrain auditory model. We thus refer to it as the midbrain Modulation Power Spectrum (mMPS).

### Distributed Stochastic Neighbor Embedding (t-SNE) of Time Averaged Mid-Level Statistics Represent Speech Articulatory Features

We then explored whether the midbrain model summary statistics account for speech articulatory features, even in its time averaged state. To do this, we projected our high dimensionality mMPS for each spoken digit in the experimental sounds, into a low dimensional space through t-distributed stochastic neighbor embedding (t-SNE) ^53^. This method projects the mMPS of each vocalization onto a 2-D space that optimally preserves the Euclidean distance between vocalizations, thus allow for visualizing the mMPS of all vocalizations in a low-dimensional space. To relate this spatial mapping to our proposed speech features, we visualized our speech measurements (F0, F1, F2) and speech groupings in the same space (Voice Group, Spoken Digit). Due to the correlation between F0 and HM (shown in Fig. 4B) we omitted HM from this analysis. Different speech groups were categorically visualized (Voice Group, Spoken Digit), while speech measurements (F0/F1/F2) were split into 20% quantiles across the whole dataset to simplify visualization.

### Generalized Perceptual Regression Modelling

Although, GLME can be used to identify factors contributing to perception, it requires categorical labels (e.g., backgrounds, words, perturbations) and articulatory measurements that are not directly represented in the auditory system. Here we use *Generalized Perceptual Regression*^2,43^, a modelling framework that links dimensionality reduced neural summary statistics of sounds to perceptual outcomes. Using a sound computable auditory model, the mMPS is used as a biologically plausible feature set to predict human perceptual trends. These auditory model features (mMPS) are then mapped to behavioral responses with a logistic regression output model. Thus, GPR combines an auditory model front end with a logistic regression output to produce a quasi-linear mapping between sound summary statistics and behavioral responses. The GPR model is fit to behavioral measurements using Bayesian optimization and validated using a 25-fold cross validation. The model was fit for each individual participant, using the 9,900 single digit responses, and mMPS features for both the foreground and background.

Given the high dimensionality of the mMPS feature space (82 frequencies x 38 temporal modulations x 8 spectral modulations = 24,928), we first performed dimensionality reduction to minimize the possibility of overfitting. Principal component analysis was applied to each training set fold during optimization and individually for each mMPS (Foreground, Background, Interaction):

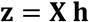

where

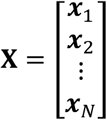

is a *N* x *M* matrix ( *N* training sounds, *M*=24,928 features) containing the sound features, ***x****_n_* (1 x *M*), **z** (*N* x *L*) is a matrix containing the dimensionality reduces sound features (principal component scores), and **h** (*M* x *L*) is a whitening matrix containing the top *L*=15 singular values of the covariance matrix **X***^T^***X** and which transforms the sound feature set from *M* (24,928 features) to *L* (15 features) dimensions. This procedure was applied separately for the foreground (*f*), background (*b*) and interaction (*i*) mMPS terms and each component was combined to generate

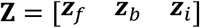

a dimensionality reduced input feature matrix (*N* x 45) that is used to predict behavioral responses.

Predictions for single sound trials are performed with a logistic regression model that maps the dimensionality reduced features (**Z**) to probabilities for performing correct decisions. The predicted outputs are obtained by first predicting the log-odds ratio:

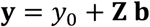

where

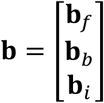

and where the **b***_f_*, **b***_b_* and **b***_i_* are each column vectors (*L* x 1) containing the logistic regression weights associated with each of the three input features (foreground, background, interaction). Finally, the predicted output probabilities of performing a correct decision are obtained by applying a logistic nonlinearity to the log-odds ratio:

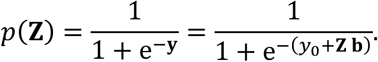

The model can also be fit with ancillary regressors and, here, we paired the training features with the serial position of the digit (1, 2 or 3) to account for any potential ordering effects.

Finally, we note that the regression weight vectors, **b***_f_*, **b***_b_* and **b***_i_*, represent regression model weights that map biologically plausible dimensionality-reduced sound representations to the log-odds ratio, and thus the probability of performing correct behavioral decisions. Thus, they can be thought of as *perceptual transfer functions* that represent the perceptual sensitivity and map the dimensionality reduced sound features to correct decisions. These regression weights can be projected back onto the original midbrain model space to allow for ease of visualization:

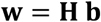

where

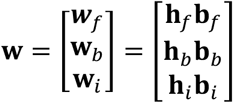

contain the vectorized perceptual transfer functions in the original auditory model space and where

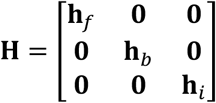

is a diagonal matrix containing the principal component whitening matrices for each feature representation (foreground, background, interaction). Thus, the individual weighting vectors can be re-ordered and visualized in the original midbrain model space (*F_l_* vs. *f_m_* vs. Ω*_n_*) which provides an interpretable account of how features map onto perception. Throughout, we refer to the GPR model weights **w** as *perceptual transfer functions* since they provide an interpretable account of each participant’s perceptual sensitivity to different acoustic features as represented in the original auditory model space.

### Reduced Feature Models

To explore what features may be driving variability in accuracy in both the voice and digit projections, we compared the performance of the full GPR model with several reduced models. Specifically, we varied the stimulus task features used in the model (foreground, background, foreground-background interaction) and varied the stimulus representation (spectral features alone or including spectral and temporal modulations). We fit GPR to each reduced feature set, using the method described above. Here, to prevent overfitting and ensure an equitable comparison across models, each feature set was dimensionality reduced to 15 PCs. We then evaluated the k-fold cross validated prediction sets scored according to the R^2^ for the voice and digit projection average accuracy, as well as single trial accuracy. Single-trial accuracy was assessed using receiver operating characteristic (ROC) relative to the single-digit behavioral responses, resulting in a summary area under the curve (AUC) metric for each participant. Separate from the averaged VP and DP scores, the AUC allows us to observe fluctuations in performance for single trials. Thus, the AUC could be helpful in exploring the effects of individual trial differences that could be accounted by the local mMPS statistics. Reduced models were compared for each of the different model metrics, reduced model comparisons for the DP and VP were done through a paired fisher z-transform test, while the AUC was compared via paired t-test. These statistical analyses were extended to both the population model and glimpsing/static model (see below).

### Individual vs Population Models to Predict Within and Across-Participant Behavior

While the perceptual transfer functions have the advantage that they reflect single-participant behavior, we also sought to test whether a single population-level model could generalize and predict performance on unseen participants. Thus, we altered our GPR method to do leave-one-out cross validation across participants. We sub-selected an individual K-fold from the original models per participant and averaged across 9 participants to predict the remaining, held-out participant. We then compared the performance of these “population models” to the “individual models” described above.

### Comparing Glimpsing vs Static Background Models

Glimpsing, the idea that soft sound segments or silent gaps in a background sound produce local increases in SNR, has been proposed as one mechanism for segregating speech from noise ^14,15^ . Since our model estimates the local mMPS summary statistics overlapping each digit, it has access to stimulus power of the speech and background at a given time point during the digit duration. Thus, in its original form, the model could potentially account for “classical glimpsing”. To test whether glimpsing can be measured with the GPR model, we compared the above described GPR model (referred to here as the *glimpsing model*) to a static GPR model variant which uses the time-average mMPS over the background segments that overlap the three digits in each sound to make accuracy predictions (the *static model*). As with the reduced, and population models, we assessed the performance by calculating the variance explained for the VP and DP averages, in addition to calculating the AUC for both the static and glimpsing models. While the VP and DP variance explained could be less susceptible to single trial variance, the AUC represents single trial performance. To explore if certain backgrounds were prone to glimpsing, and whether specific background features contributing to glimpsing could be identified, we compared the AUC from the glimpsing and static models for each background (OR, SM and MM). Here positive AUC difference suggests that the background is sound is prone to glimpsing, while a negative difference suggests that time-averaged summary statistics better describe perception ^54^.

## Funding

This work was supported by the National Institute on Deafness and Other Communication Disorders of the National Institutes of Health under awards R01DC020097 and R01DC023478. The content is solely the responsibility of the authors and does not necessarily represent the official views of the NIH. The funders had no role in study design, data collection and analysis, decision to publish, or preparation of the manuscript.

## Notes

### Competing Interest Statement

The authors have two patent pending related to methods used in the study.

### Summary of Updates

Slight modifications to figures and modifications to text to improve clarity.

